# Biosensor monitoring of naphthenic acids remediation in mesocosms and constructed wetlands: a head-to-head comparison with orbitrap mass spectrometry

**DOI:** 10.64898/2026.06.02.729691

**Authors:** Tyson Bookout, Ian J Vander Meulen, Amy-Lynne Balabera, Dani Degenhardt, John V Headley, Shawn Lewenza

## Abstract

Oil sands process-affected water (OSPW) contains complex mixtures of naphthenic acids (NA) that are the central targets for water treatment and reclamation. Here, we compared three whole-cell bacterial NA biosensors with Orbitrap mass spectrometry (MS) for quantifying NA remediation in greenhouse mesocosms and a pilot-scale constructed wetland. Solid-phase extracts from both systems were analyzed in parallel by Orbitrap MS and biosensor assays, enabling direct comparison of biosensor-derived NA estimates with MS-derived naphthenic acid fraction compounds (NAFC) concentrations. Across both treatment systems, biosensor outputs broadly tracked declines in NAFC measured by Orbitrap MS, and positive linear relationships were observed between methods. Biosensor 2 (3680-lux) and biosensor 3 (atuA-lux) showed, early rapid decreases in NA, whereas biosensor 1 (marR-lux) frequently remained elevated later in treatment. These differences are consistent with the distinct chemical response profiles of the biosensor panel and suggest that biosensor outputs reflect shifts in NA mixture composition as remediation proceeds. This interpretation is supported by the published Orbitrap analysis of the constructed wetland, which showed decreasing O_2_-NAFCs and increasing O_3_/O_4_-containing species, consistent with oxidative degradation. Together, these results support the use of NA-responsive biosensors as rapid and scalable complementary tools for tracking remediation trends, while Orbitrap MS remains the reference method for molecular-level characterization.

## Introduction

Oil sands surface mining in northern Alberta generates very large volumes of oil sands process-affected water (OSPW), a complex industrial wastewater containing inorganic clays and suspended solids, residual hydrocarbons, with a major fraction of the organic compounds consisting of naphthenic acids (NAs) (1, 2). The formula C_n_H_2n-z_O_2_ is used to describe classic NAs, where n is the number of carbons and z indicates the number of hydrogens lost due to ring formation (3, 4). NAs are complex mixtures of monocyclic, polycyclic, and acyclic alkyl-substituted carboxylic acids, and naphthenic acid fraction compounds (NAFCs) include a broader range of compounds based on carbon number, double bond equivalents, number or rings, aromatic rings or heteroatom content (3–5). Due to the accumulation, persistence and toxicity of NAs (6), they are the central target for water treatment strategies intended to enable long-term reclamation and, ultimately, controlled release of legacy process waters. As regulatory and industrial frameworks increasingly emphasize treatment of stored tailings water, there is growing demand for monitoring approaches that can track NA removal and transformation during remediation at operationally relevant scales.

Constructed wetlands have emerged as a promising nature-based solution for NA attenuation (7, 8). Field and pilot-scale studies indicate that wetland plants and their associated microbial communities can contribute to NA removal through biodegradation and compositional transformation processes. Mesocosm systems provide a simpler experimental wetland model to further show that treatment performance can vary across conditions and designs, and that NA remediation often involves changes in NA molecular composition, in addition to changes in total concentration (9–11). Despite these advances, monitoring tools that are sufficiently specific, rapid, cost-effective, and scalable for high-frequency screening and operational decision-making remain as a practical barrier for large-scale wetland implementation.

At present, high resolution mass spectrometry (HRMS), including Orbitrap analysis, represents a gold standard for NAFC characterization (12, 4, 5). HRMS provides deep molecular-level resolution, enabling assignment of hundreds of NAFC formulae and evaluation of carbon number distributions, double bond equivalents, heteroatom classes, and oxidation trends. This level of detail is important for evaluating wetland treatment performance and documenting NAFC compositional shifts during attenuation. HRMS workflows typically require organic solvent or solid phase extraction to concentrate naphthenic acids, expensive, specialist instrumentation, and complex analysis, which may limit cost effective and high-throughput monitoring programs.

Whole-cell bacterial biosensors provide a complementary analytical approach for NA detection and quantification. These systems exploit the natural sensing and regulatory responses of bacteria that detect numerous organic compounds, and produce a simple, measurable output (e.g., bioluminescence) (13, 14). Biosensors can offer major advantages for environmental monitoring, including rapid assay time, low per-sample cost, minimal instrumentation, and high-throughput compatibility. Recently, we developed a panel of NA-responsive biosensors in a tailings pond-derived *Pseudomonas* strain using transcriptomic identification of NA-responsive genes, generating promoter-reporter constructs with distinct, and concentration-dependent responsiveness to different NA subgroups (15).

Here, we present a head-to-head comparison of a three biosensor panel to Orbitrap HRMS studies across two OSPW remediation contexts: (i) a greenhouse mesocosm treatment model, representing a controlled system in which soil- and plant-driven effects can be assessed more directly (11); and (ii) a constructed wetland treatment system at the Kearl site, representing a field-relevant and spatially heterogeneous system (8). Biosensor methods were developed for rapid, qualitative results within 24 hours, or semi-quantitative NA analysis providing a mg/L estimate, enabling a direct comparison across analytical platforms. Here we demonstrate the potential for biosensor-based monitoring as a rapid, simple, and scalable complement to HRMS for NA remediation assessment.

## Materials and methods

### OSPW Treatment Systems and Water Sampling

A controlled greenhouse mesocosm system was used as a small-scale, constructed wetland analog to evaluate NAFC attenuation under defined treatment conditions, as has been previously reported elsewhere (11). Mesocosms were operated as closed-loop, surface-flow treatment systems containing wetland substrates and OSPW, enabling direct comparison among treatments that differed by the presence/absence of soil and plants. Treatments included an untreated water-only condition, a soil-amended control, and planted treatments incorporating the wetland plants *Carex aquatilis* and *Typha latifolia*. Mesocosms were run as recirculating units over a multi-week experimental period with repeated sampling across a time course to quantify changes in NAFC concentration during treatment. The overall mesocosm construction and operational parameters follow previously described wetland mesocosm approaches for OSPW remediation studies (9–11).

Pilot-scale NAFC remediation was previously reported using the Kearl Treatment Wetland, an engineered constructed wetland treatment system operated as a contained, recirculating free water surface wetland with both shallow vegetated and deeper open-water segments (8). During operation, OSPW was introduced at the forebay and circulated end-to-end through the wetland under a targeted, full-volume recirculation time of ~14 d, enabling evaluation of NAFC concentration trends over the growing season. Water samples were collected across multiple wetland sub-areas representing spatial heterogeneity in depth and vegetation (e.g., forebay, shallow areas, deep pools) and over repeated time points across a seasonal treatment timeline.

### Solid phase extraction (SPE) to prepare naphthenic acid fraction compounds from OSPW

Water samples from mesocosm and wetland cells were collected into pre-cleaned amber glass bottles and stored cold prior to shipment and processing. Samples were extracted using solid-phase extraction (SPE) to recover acid-extractable organic fractions containing NAFCs, as previously described (5). Briefly, Biotage ENV+ SPE cartridges (6 mL / 200 mg) were rinsed with MilliQ water (≥ 18.2 MΩ; 6 mL), LCMS-grade methanol (Fisher Scientific Canada), and reconditioned with 6-12 mL of MilliQ water. Aliquots of samples (~100 mL; exact volume recorded) were acidified (pH < 2). Following conditioning, samples were drawn through cartridges at 3-5 mL/min, washed/desalted with a further 6 mL of MilliQ water, dried under gentle vacuum, then NAFCs were subsequently eluted in methanol, dried under nitrogen, and reconstituted in 50:50 water:acetonitrile (LCMS-grade; Fisher Scientific Canada) with 0.1 % NH_4_OH (28 – 30 %). The same SPE-derived extracts generated for Orbitrap HRMS analysis were subsequently used for biosensor testing in the present work, enabling direct head-to-head comparison between Orbitrap-based NAFC quantification and biosensor-derived concentration equivalents using identical extracted organic material.

### Quantitative detection of luminescence from biosensors to estimate naphthenic acid concentration in OSPW

For quantitative NA detection, solid phase extraction was used to concentrate the organic compounds in OSPW by 100-fold, as described above. Five µl (50:50 acetonitrile:water with 0.1% NH_4_OH)) of NAFC extracts were resuspended in 94 µL of Basal minimal medium (BM2) media (0.5 mM Mg^2+^, 20 mM succinate), which were inoculated with 1 µL of previously characterized strains: biosensor 1 (marR^H^::lux), biosensor 2 (3680^2^::lux) or biosensor 3 (atuA^wt^-lux) (15). The *lux* gene expression assays were conducted in liquid cultures grown in black 96-well, clear bottom plates (Thermo Scientific). A Breathe-Easy® (Sigma-Aldrich) membrane was used to prevent evaporation during a 15-hour protocol in a PerkinElmer 1420 multilabel counter Victor^3^. The plate reader protocol (2 sec shake; luminescence in counts per second (CPS), growth (OD_600_) included 45 time points, taken every 20 minutes. Bioluminescence/gene expression (CPS) was normalized by dividing the CPS by the growth (OD_600_) for each read. When NA exposure caused a gene induction, this is interpreted as detection of NAs. The fold induction was calculated by dividing the CPS/OD_600_ values of the biosensor OSPW response by the control condition without NAs at each time point. We plotted data from the time point that caused maximum induction.

### Biosensor standard curves for total NA estimation

Standard curves for each biosensor were obtained by measuring luminescence as described above, where each strain was exposed to a range of NAFC concentrations between 160-440 mg/L, using a representative NAFC extract from industry water. Since NAFCs in extracts were tested at 10X their concentration in OSPW, this range corresponds to 16-44 mg/L in water. The linear regression to gene expression was determined and represented as the following equations: biosensor 1 (marR-lux), Y = 0.02254*X - 2.647 (R^2^ = 0.9849); biosensor 2 (3680-lux), Y = 0.01033*X - 0.06935 (R^2^ = 0.9701); biosensor 3 (atuA-lux); Y = 0.009495*X + 0.1472 (R^2^ = 0.9591). We measured the biosensor responses to extracts from mesocosm or wetland treated water samples to determine the fold induction and then used the luminescence standard curves to extrapolate and determine a mg/L value.

### Linear regression analysis of biosensor and Orbitrap measurements

To assess agreement between biosensor-based estimates of total NA and Orbitrap-derived measurements from the same SPE extracts, ordinary least squares (OLS) linear regression was performed separately for each biosensor and treatment condition. For mesocosm samples, regressions were conducted within each treatment group across the time series. For constructed wetland samples, regressions were conducted within each wetland compartment across sampling dates. In each analysis, the Orbitrap-derived value was treated as the independent variable (x) and the biosensor-derived NA estimate as the dependent variable (y). The strength of the linear association was evaluated using the coefficient of determination (R^2^), and the fitted slope and intercept were used to assess proportional agreement between methods. Regression plots were generated for visualization of method-to-method relationships.

### Qualitative biosensor detection of naphthenic acids in water samples

To minimize the processing of water samples, raw mesocosm or wetland samples (2-4 ml) were evaporated to dryness by rotary evaporation at 60 ºC and resuspended in 25 µL of double distilled H_2_O. Biosensor strains were grown in overnight cultures, diluted 1/10 and normalized to the same optical density (OD_600_ = 0.5), and streaked as a lawn onto dilute 1/32 LB agar plates. Dilute LB mimics minimal media and limits bacterial growth for optimal luminescence responses. Four µL of each concentrated water sample was spotted in a grid pattern onto agar plates, which were incubated overnight at 30 ºC, and then imaged using the Chemi-Doc bioluminescence imaging system (BioRad). Plates were inspected for luminescence ring patterns from all samples in this qualitative ‘agar spot’ assay.

### Composite Biosensor Index Calculation

To provide an integrated measure of biosensor response for the panel of three biosensors, a composite biosensor index (CBI) was calculated. The CBI values for the wetland system were calculated from two independent biological experiments, each comprising three technical replicates per biosensor at each sampling point. For each biosensor and time point, NA concentration estimates (mg/L) were normalized to the initial time point (t_0_) within each wetland compartment. The normalized responses of the three biosensors were then averaged to generate the composite index according to the equation: CBI = 1/3 (B_1_^norm^(t) + B_2_^norm^(t) + B_3_^norm^(t)).

## RESULTS

### Quantitative monitoring of NA remediation using mesocosm and constructed wetlands systems designed for treating oil sands process-affected water

To enable direct comparison between analytical platforms, all water samples collected from the mesocosm, and constructed wetland treatment systems were first processed by solid-phase extraction (SPE) to concentrate the naphthenic acid fraction compounds (NAFC). The same SPE extracts were then used for both Orbitrap mass spectrometry (MS) analysis and biosensor-based quantitation of total naphthenic acids (NA). The naphthenic acid biosensor strains were previously described and shown to detect and respond to complex NAFC mixtures by producing a quantitative bioluminescence signal in proportion to the NAFC concentration (15). The specificity of the biosensors was also tested with a limited set of available NA compounds, and each biosensor was shown to detect a distinct, minimally overlapping subset of model NA structures. The profile of model compounds detected by each biosensor provide a guideline to determine what structural features are likely detected when exposed to complex NAFC mixtures (15).

Biosensor 1 (marR-lux) detects classic, single-ring O_2_ compounds (cyclohexane acetic acid, CHAA) and other complex NA structures, including the pyridine ring-containing compound fusaric acid, 5,6,7,8-tetrahydro-2-naphthoic acid (THAN; aromatic O_2_), 3-phenylpropionic acid (PPA; aromatic O_2_), cyclohexyl succinic acid (CHSA, O_4_) and adamantane carboxylic acid (ACA, diamondoid O_2_) (15). Biosensor 2 (3680-lux) detects mostly classic, single ring O_2_ (NA) compounds such as cyclopentane carboxylic acid (CPCA), cyclohexane carboxylic acid (CHCA), 3-phenylpropionic acid (PPA) and cyclohexane hexanoic acid (CHAA) (15). Biosensor 3 (atuA-lux) is a fatty acid biosensor, that responds mainly to medium length, linear and branched fatty acids (C6-C18), but also to commercial mixtures of naphthenic acids that contain a majority of Z=0, acyclic fatty acid compounds (15). Fatty acids (Z=0) are commonly found in OSPW and are likely degradation products of naphthenic acids.

In the mesocosm treatment system, the time-series data showed that both Orbitrap MS and the biosensors were able to quantitatively demonstrate remediation trends where NAFC concentrations decreased in all treatments, over time (Fig 1A). The Orbitrap MS profiles showed gradual reductions in total extractable NAFCs over time, whereas the biosensors often showed stronger or earlier decreases, particularly in the planted treatments. In contrast, the no-treatment control (T1) showed comparatively flat MS profiles over the course of the experiment, consistent with persistence of the total NA pool in the absence of active remediation (Fig 1A). In addition, we calculated the area under the curves (AUC) in the mesocosm system as an approach to further estimate the total NAFC concentrations during treatment. In Figure 1C, it is shown that the AUC values in the non-treated controls are higher than the treatment conditions, which is generally true for all methods. MS estimates for NAFC concentrations showed a highly significant decreases during treatments, and among the biosensors, marR also showed a significant overall decrease during treatments, whereas 3680 and atuA showed the same downward trend but did not reach significance across the treatment period (Fig 1C). According to the MS results, the T5 treatment with both plants caused the maximum reduction in NAFCs, followed by T3 and T4, each plant individually (11) (Fig 1A). Biosensors indicated that planted systems were more effective than soil, but there was no consensus as to the best plant treatment (Fig 1A).

**Figure 1.**
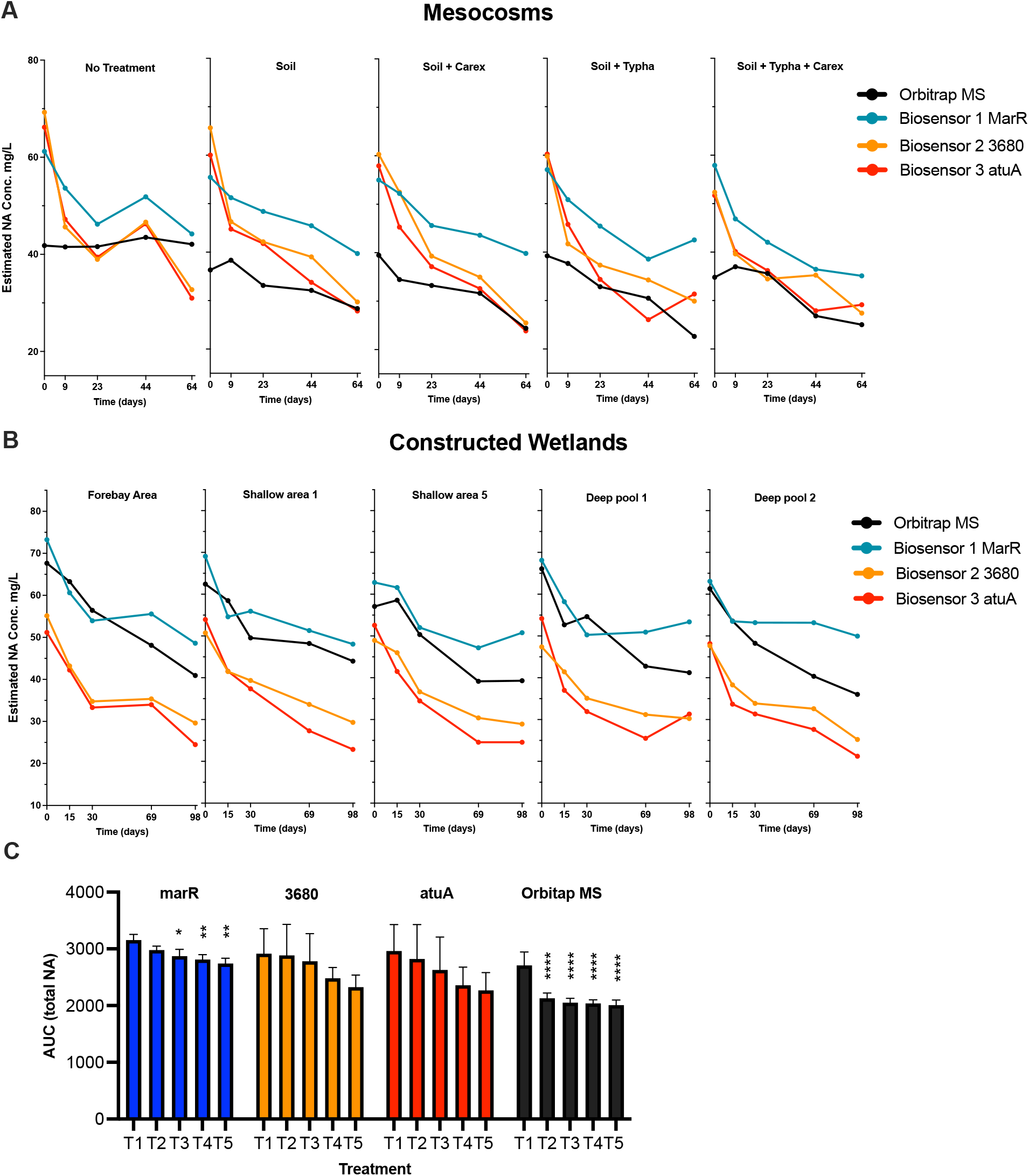
Time series of NA degradation in mesocosms and constructed wetlands as measured by biosensors and orbitrap MS. Solid phase extraction was used to concentrate NAFC in all water samples and were tested in parallel with biosensors and Orbitrap MS in **A)** 5 mesocosm conditions and **B)** different sections of constructed wetlands over time. Total NAFC (mg/L) was determined by orbitrap MS (black line) and total NAs with three independent biosensors (blue, red, orange lines), which produce luminescence in proportion to the amount of NA in the water. Biosensor values shown are the averages of 6 technical replicates, from 2 independent SPE extracts, and MS values for mesocosm and wetland extracts are the average of 4 or 2 independent SPE extracts, respectively. **C)** Area under the curve (AUC) from the mesocosm (T1, no treatment; T2, soil; T3, soil + Carex; T4, soil + Typha; and T5, soil + both plants) and wetland data in panel A, B. Values shown are the averages of 9 technical replicates, from 3 independent SPE extracts. A significant reduction in NA was determined by a one-way ANOVA to test for an overall treatment effect, followed by Dunnett’s multiple-comparisons test to compare each treatment with the control. Significant NA reduction relative to the control is shown with asterisks (* p < 0.05, ** p < 0.01, **** p < 0.0001).

A similar overall pattern was observed in the constructed wetland treatment system, where both Orbitrap MS and the biosensors tracked declining NAFC and NA concentrations across the five monitored wetland compartments (Fig 1C). The forebay (FA) and the two shallow areas (SA1 and SA5) showed progressive decreases in NAFC and NA concentrations over time by both MS and biosensor readouts, consistent with attenuation during passage through the wetland. The deep pools (DP1 and DP2) also showed declines, although these compartments retained higher concentrations for longer periods in some methods, suggesting slower removal or persistence of more resistant NAFC and NA fractions.

Within the mesocosm and wetland time-series, biosensor 2 (3680-lux) and biosensor 3 (atuA-lux) frequently showed early, rapid and sustained declines, whereas the marR-lux signal often declined less sharply, and in several cases appeared to flatten at later time points (Fig 1). This pattern is consistent with the known response profiles of these biosensors. The marR-lux sensor responds not only to certain classic O_2_ NA compounds but also to more recalcitrant compounds (aromatic, diamondoid, multi-ring, O_4_,) (15). As remediation proceeds, the 3680-lux and atuA-lux biosensors indicate that the more easily degraded, single ring and acyclic O_2_ compounds are preferentially transformed, and the remaining NAFC pool may become proportionally enriched in recalcitrant compounds. This pattern would result in a higher marR-lux biosensor response at later time points, while the 3680-lux and atuA-lux biosensors show stronger declines throughout. These biosensor results align with the molecular analysis of NAFCs by Orbitrap MS, where O_2_-NAFCs were decreasing over time, along with an increasing proportion of O_3_ and O_4_-NAFCs, which is indicative of oxidative degradation (8). In summary, the biosensors tracked the NA remediation trends shown with Orbitrap MS and also provide general insights towards the kinds of NAFCs that are being preferentially degraded.

### Biosensor estimates of total NA concentrations positively correlate with values from Orbitrap MS

To determine the correlation between biosensor-derived estimates of total NA and Orbitrap MS NAFC measurements, we performed ordinary least squares (OLS) linear regression analysis for each biosensor within each mesocosm treatment and wetland compartment. The strength of the association was quantified using the coefficient of determination (R^2^), and the regression slopes were used to assess proportional agreement between analytical approaches.

In the mesocosm system, biosensor-derived NA estimates were positively associated with Orbitrap MS NAFC measurements across the remediation treatments (Fig. 2A). Correlations were strongest in the treated mesocosms, compared to the untreated control where MS-measured concentrations did not change substantially over time and did not provide a meaningful dynamic range for regression (data not shown). All three biosensors showed positive, treatment-specific relationships with MS, although the strength of correlation varied among biosensors and among treatment conditions. Overall, biosensor 1 (marR-lux) tended to show the most consistent agreement with MS across the mesocosm treatments, whereas biosensors 2 (3680-lux) and 3 (atuA-lux) showed more variable agreement across treatments (Fig 2A).

**Figure 2.**
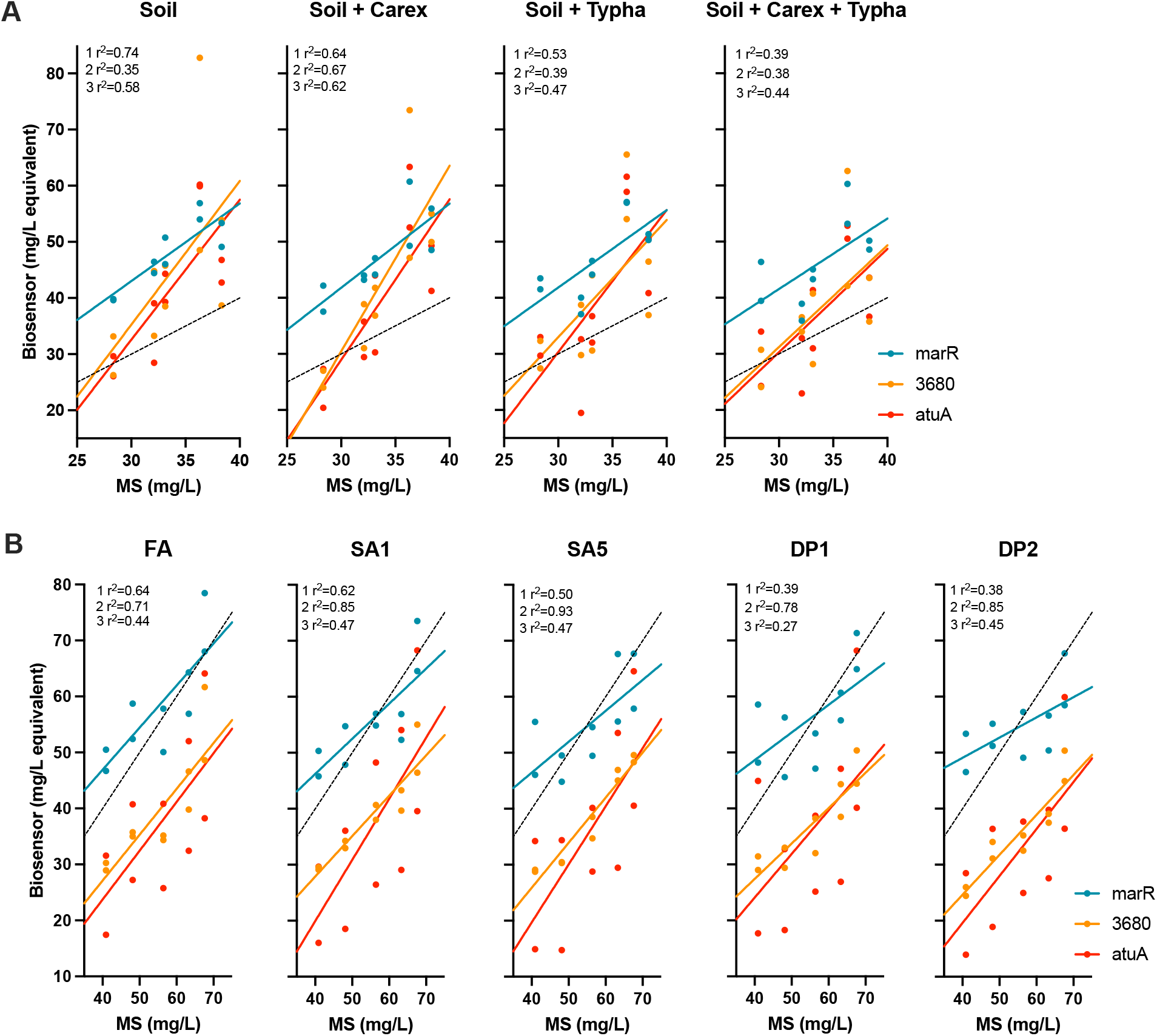
Ordinary least squares (OLS) regression analysis comparing biosensor-derived estimates of total NA with Orbitrap MS measurements. Treatment of OSPW in (A) mesocosm and (B) constructed wetlands. For each panel, Biosensors 1 (marR-lux), 2 (3680-lux), and 3 (atuA-lux) were plotted against Orbitrap MS values obtained from the same SPE-derived extracts. Solid coloured lines indicate OLS regression fits for each biosensor, and the dashed line indicates the 1:1 relationship. R^2^ values for each biosensor are shown within each panel. Positive relationships between biosensor output and Orbitrap MS indicate that the biosensors broadly tracked treatment-associated changes in total NA across both remediation systems.

The wetland OLS analysis likewise showed positive linear relationships between biosensor output and Orbitrap MS measurements across all wetland compartments (Fig. 2B). As in the mesocosms, correlation strength varied by biosensor and by sampling location. However, the overall pattern in the wetland was somewhat different, with biosensor 2 (3680-lux) generally showing the strongest and most consistent agreement with Orbitrap MS across compartments, while biosensors 1 and 3 showed more variable performance depending on location. The regression lines were also commonly offset from the 1:1 reference line, indicating that biosensor-derived mg/L-equivalent concentrations did not show strict equivalence with MS-derived concentrations. Although the relative agreement of individual biosensors differed between the mesocosm and wetland systems, all three biosensors broadly tracked the same treatment-associated concentration changes measured by Orbitrap MS.

### Comparison of NA remediation rates within the early stages of treatment

We analysed the rate of declines (slope) during the early stages of treatment and showed that NA concentrations generally declined across all mesocosm and wetland sections, although the magnitude of decline differed among methods (Fig 3). In the mesocosm and several sections of the wetland, Biosensor 2 (3680-lux) and Biosensor 3 (atuA-lux) exhibited steeper negative slopes than Orbitrap MS, indicating more rapid loss of the NA fractions detected by these biosensors than of the total extractable NAFC pool measured by MS (Fig 3). By comparison, the marR-lux biosensor often showed shallower declines at early remediation stages (Fig 3), in addition to the flat rates of NA remediation during later time points (Fig 1), consistent with interpretation that the compounds sensed by marR are more difficult to degrade. Together, these results suggest that the biosensors provide complementary kinetic information and may capture early changes in bioavailable NAFC fractions during wetland treatment. Where MS methods detect the total number of charged, organic and ionized compounds, the biosensors only detect compounds that are bioavailable, that enter the cell and may be degraded. MS provides high resolution information on NAFC transformation during remediation, and the biosensor readout is a lower resolution, kinetic indicator of rapid degradation of compounds preferentially degraded.

**Figure 3.**
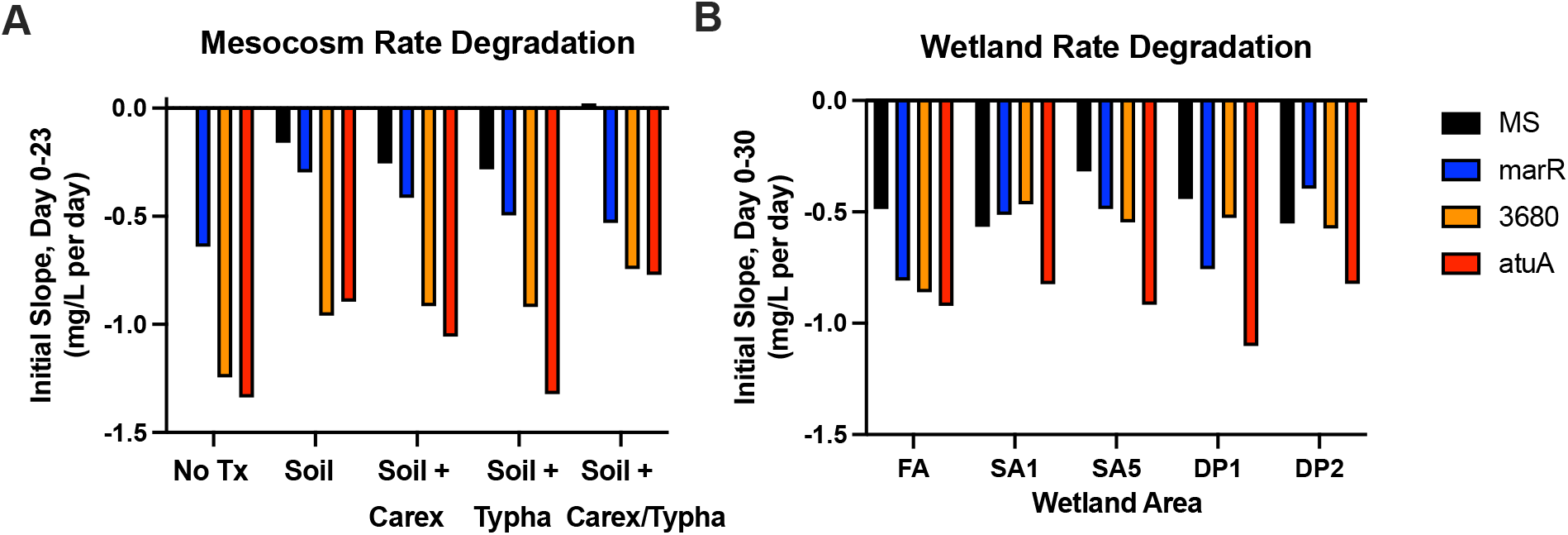
NA degradation rates during the early time frame of remediation. The rates of remediation were calculated during **A)** the first 23 days of mesocosms experiments or **B)** the first 30 days of wetland experiments, for all treatments, wetland areas and all monitoring methods. Negative values indicate decreasing rates of NA remediation.

### Rapid, qualitative biosensor monitoring of NAs in mesocosm and constructed wetland remediation systems for treating oil sands process-affected water

We aimed to design a simple method to utilize naphthenic acids biosensors to provide a rapid, output and indicator of NA remediation. For this experiment, small volumes (2-4 ml sample) of water samples were fully evaporated in order to resuspend the resulting pellet in 25 µl water, effectively concentrating the water 80 to 160-fold. The bacterial biosensor strains were streaked as a consistent lawn on the surface of agar plates, and a small volume (4 µl) of concentrated water was spotted onto agar plates. Dilute LB growth medium (1/32) mimics minimal media and limits bacterial growth for optimal luminescence responses. After an overnight incubation, the plates were imaged for the detection of luminescent rings or haloes, that form directly where water samples were spotted.

Figure 4A depicts the luminescence biosensor responses on agar plates when exposed to all mesocosm water samples from a time series of OSPW treatment in greenhouse mesocosms (0-64 days). When exposed to untreated or early stage treated water, all three strains produce a luminescent ring at the edge of a zone of killing, the latter indicates that the concentrated water is toxic to the biosensors. Agar experiments allow diffusion of compounds in samples, where NA can diffuse away from the initial spot, producing an NA gradient and results in a ring of luminescence around the killing zone (Fig 4). In the top row, exposure to untreated water produced strong biosensor responses at all time points during the 64-day treatment period in mesocosms, where the luminescence ring surrounds a zone of killing (Fig 4A). Using quantitative NA data generated from Orbitrap MS (Fig 1), it is known that the total NAFC concentrations in all untreated water samples were between 39-47 mg/L (8).

**Figure 4.**
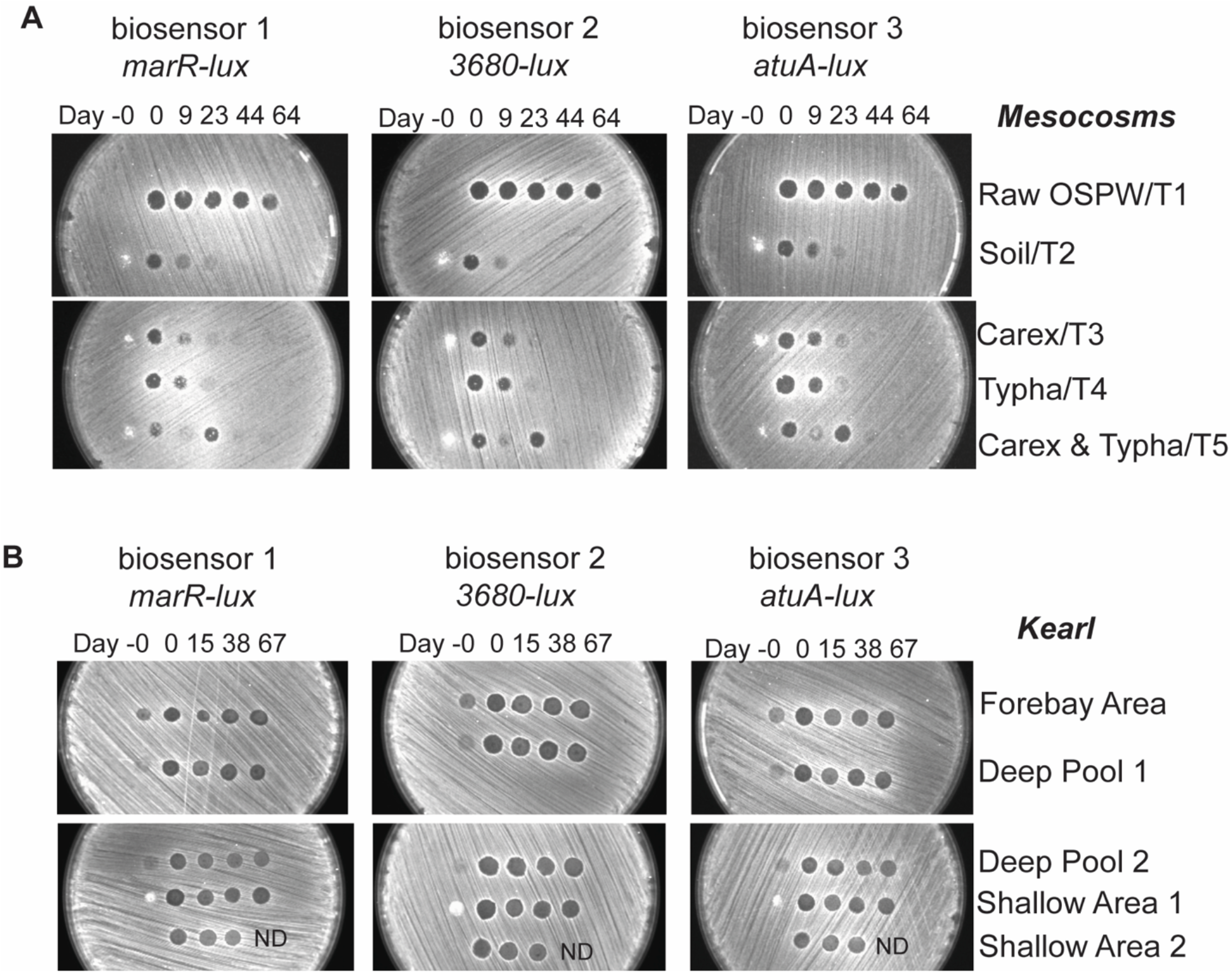
Qualitative, rapid NA detection of NA with mesocosms and wetland water samples spotted onto agar plate. All water samples from the **A)** mesocosm experiment and **B)** constructed wetland treatments were concentrated 80-fold and spotted onto a lawn of bacterial biosensor. After incubation, luminescent rings were observed around the spotted samples. A time series including before the addition of OSPW (day −0), and then days 0-64 or 0-67 of treatment in mesocosms and wetlands, respectively. These are images from one of four replicates, and the patterns were consistent between all replicates. ND, not determined.

All four treatments in mesocosm experiments were shown to be effective in reducing NA concentrations, as observed by the gradual disappearance of the luminescence between day 0 and day 23 (Fig 4A). The zone of killing also disappears, which indicates that NA toxicity is being ameliorated, leading to less bacterial killing, and less luminescence (Fig 4A). According to Orbitrap MS, all mesocosms treatments reduced the NAFC concentrations to values between 30-40 mg/L during the first 23 days (8) (Fig 1, 4). After day 44, there was no luminescence response (or killing) from all water samples tested, which indicates NA remediation below the level of detection for this bioassay (Fig 4).

Figure 4B depicts the luminescence biosensor responses on agar plates when exposed to all constructed wetland samples from a time series of OSPW treatment (0-67 days) (8). The biosensors indicated a strong luminescent ring response across the entire 67-day period, and in all cells of the wetland that were tested (Fig 4B). The zones of luminescence at the edge of the zone of killing for wetland samples were similar to the responses to untreated water in the mesocosm study (Fig 4A). According to the orbitrap MS of these wetland water samples, the total NA concentrations ranged from roughly 48 - 70 mg/L (8) (Fig 1). Since NA concentrations remained above 40 mg/L during treatment, the agar spot method could not elucidate efficacy differences across wetland treatment areas (Fig 4B). However, this method using minimally treated water samples provided a rapid and simple luminescence ring indication of high NA concentrations above 40 mg/L, which gradually weakens in response to mid-range NA concentrations (30-40 mg/L), and there is no response below 30 mg/L.

### Composite biosensor index effectively demonstrates NA remediation trends

In addition to examining the responses of three independent biosensors, we calculated a composite biosensor index (CBI), defined as the mean relative change in biosensor-derived concentration values over time. The index reflects integrated trends in the remediation of multiple NA subclasses, including compounds differing in structure, recalcitrance, and bioavailability. Across most wetland compartments, the CBI exhibited a progressive decline over time, consistent with transformation of NA during treatment, followed by a plateau in later sampling stages (Fig. 5A). The reduced rate of change observed at later time points corresponded to the behavior of the marR-based biosensor, which showed comparatively slower declines under these conditions (Fig. 1). Collectively, the CBI index provides a simple representation of integrated biosensor responses and may be useful for summarizing remediation trends during treatment.

**Figure 5.**
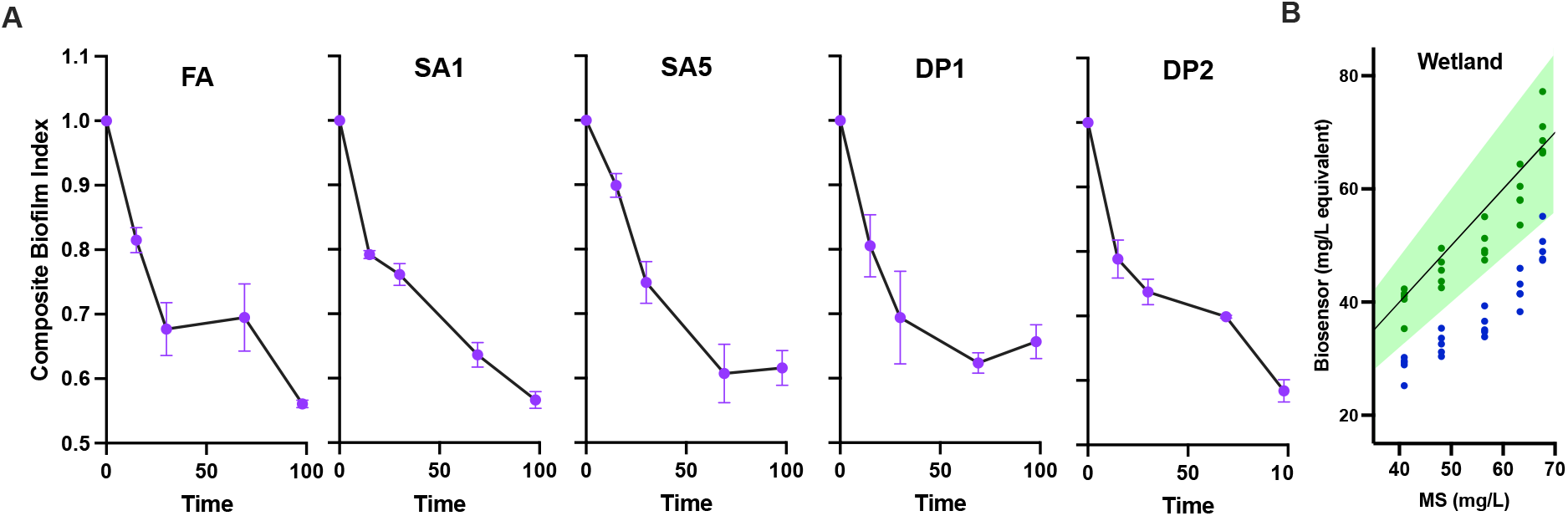
Composite biosensor index as an integrated measure of treatment-associated trends over time. **A)** Time-series plots of the composite biosensor index (CBI) for the five monitored constructed wetland compartments: forebay (FA), shallow area 1 (SA1), shallow area 5 (SA5), deep pool 1 (DP1), and deep pool 2 (DP2). The CBI values were calculated separately for 2 biological replicates each averaging 3 technical replicates, which were normalized to the initial time point and averaged across the three biosensors. **B)** Pairwise plot comparing 3680 biosensor-derived values to Orbitrap MS measurements for all forebay area wetland samples. The green region represents the mean Orbitrap MS values for NAFC concentration ± 20%. Raw biosensor 2 (3680-lux) values (blue) sit mostly below the target range, and converted biosensor values (black) align within the gold standard reference range after applying a multiplication factor of 1.4.

To further evaluate the practical utility of biosensor-based measurements, we applied a correction factor adjustment of biosensor data, in order to bring biosensor-derived concentrations within an acceptable ± 20% range of the mean target values generated by Orbitrap MS. Using the biosensor with the strongest correlation to MS in the wetland system (Biosensor 2, R^2^ = 0.78), the raw biosensor 2 (3680-lux) data fell outside of the green target area, and it generally underestimated the total NA concentrations (Fig 5B). Biosensors are not expected to provide direct equivalence to MS measurements due to many methodological differences, however, a simple multiplication factor of 1.4 was applied to align biosensor-derived values with MS-derived concentration ranges (Fig 5B). Importantly, this approach does not imply analytical equivalence but rather provides a practical means of calibrating biosensor outputs to the gold standard method for operational monitoring purposes.

## Discussion

Across both the greenhouse mesocosms and the pilot-scale constructed wetland, the three whole-cell biosensors broadly reproduced the remediation trends measured by Orbitrap MS, supporting their value as practical tools for monitoring NAs. Positive OLS relationships between biosensor-derived estimates and Orbitrap measurements showed that all three biosensors tracked treatment-associated changes in the extracted NAFC pool, although agreement varied among biosensors and between systems. In the mesocosms, biosensor 1 (marR-lux) tended to show the strongest overall agreement with Orbitrap MS, whereas biosensor 2 (3680-lux) generally showed the most agreement across the wetland compartments. However, none of the biosensors showed a strict 1:1 relationship with Orbitrap-derived concentrations, indicating that their outputs should be interpreted as biologically informed estimates rather than direct replacements for HRMS values.

A consistent pattern across both systems was that biosensor 1 (marR-lux) generally reported the highest NA concentration estimates and often remained elevated later in treatment, whereas biosensor 2 (3680-lux) and biosensor 3 (atuA-lux) more often showed stronger early declines. This pattern is consistent with the distinct specificity profiles of the biosensor panel based on prior work with model NAs (15). Biosensor 2 is biased toward more classical, single-ring O_2_-type NA structures, while biosensor 3 behaves more as a fatty acid-responsive reporter. The rapid decline indicated by biosensors 2 and 3 is supported by the published Orbitrap MS analysis of the Kearl wetland, which showed a progressive decrease in O_2_-NAFCs together with an increase in more oxygenated O_3_ and O_4_-containing formulae, consistent with oxidative transformation (8). In contrast, biosensor 1 (marR-lux) appears to respond to a broader set of compounds that includes more persistent or compositionally complex fractions. Thus, the biosensors appear to function as proxies for total extractable NAFCs, while also providing complementary coarse compositional information about which portions of the NA mixture are being preferentially transformed during remediation.

Given the immense volume of legacy OSPW requiring treatment prior to reclamation and eventual release, there is a clear need for monitoring tools that are informative, practical, and scalable. In this context, whole-cell biosensors offer several important advantages: they are comparatively low cost, rapid, simple to perform, and capable of reliably detecting remediation-associated changes in NA concentrations across both controlled and field-relevant treatment systems. Although biosensor outputs are not equivalent to Orbitrap MS and do not provide the same molecular resolution, they consistently tracked degradation trends. If future validation confirms stable relationships with HRMS, biosensor-estimated NAFC concentrations could be calibrated with correction factors to align with expected concentration values. Accordingly, biosensors may provide a useful high-throughput screening approach for routine monitoring of NA remediation, while Orbitrap MS remains the gold standard reference method for detailed compositional characterization and confirmation.

## Acknowledgments

This work was supported from a Genome Canada Large-Scale Applied Research Project (LSARP) grant (#18207), in partnership with Genome Alberta and Genome Quebec. Co-funding was provided by the Government of Alberta through an Alberta Innovates grant and Jobs, Economy and Innovation funding.

